# k-BOOM: A Bayesian approach to ontology structure inference, with applications in disease ontology construction

**DOI:** 10.1101/048843

**Authors:** Christopher J. Mungall, Sebastian Koehler, Peter Robinson, Ian Holmes, Melissa Haendel

## Abstract

One strategy for building ontologies covering domains such as disease or anatomy is to weave together existing knowledge sources (databases, vocabularies and ontologies) into single cohesive whole. A first step in this process is to generate *mappings* between the elements of these different sources. There are a number of well-known techniques for generating mappings (also known as *ontology alignemnt*), both manual and automatic[7]. Sometimes mappings are seen as an end in themselves, with the sources remaining in a loosely connected state. However, if we want to take the next step and use the mappings to integrate the different sources into a cohesive reference ontology, then we need to translate the mappings into precise logical relationships. This will allow us to safely merge equivalent concepts, creating a unified ontology. This translation is a non-trivial step, as each mapping can be interpreted as multiple different logical relationships, with each interpretation affecting the likelihood of the others. There is a lack of automated methods to assist with this last step; this resolution is typically performed by expert ontologists.

Here we describe an ontology construction technique that takes two or more ontologies linked by hypothetical axioms, and estimates the most likely unified logical ontology. Hypothetical axioms can themselves be derived from semantically loose mappings. The method combines deductive reasoning and probabilistic inference and is called Bayesian OWL Ontology Merging (BOOM). We describe a special form k-BOOM that works by factorizing the probabilistic ontology into *k* submodules. We also briefly describe a supplemental lexical and knowledge-based technique for generating a set of hypothetical axioms from loose mappings.

We are currently using this technique to build a merged disease ontology (Monarch Disease Ontology; MonDO) that unifies a broad range of vocabularies into a consistent and coherent whole.

## 1 INTRODUCTION

Ontologies provide a cohesive representation of some knowledge domain, such as human anatomy or human disorders. One of the characteristic features of an ontology is that relationships between elements have precise logical interpretations, usually expressed in the Web Ontology Language (OWL). For some domains, such as cellular biology, we have consensus reference ontologies such as the Gene Ontology, with broad and deep coverage of the domain. In other areas, such as diseases) there are multiple ontologies and databases with distinct perspectives and overlapping content. Here no single ontology provides the complete picture, so we would like to integrate these together into a unified cohesive whole. The most common approach here is to generate mappings connecting classes or database entities. However, mappings are only one step towards producing a unified ontology. Most mappings lack clear semantics, and so cannot be used effectively for reasoning in a description logic environment. Additionally, mappings are typically not enough to merge classes from two ontologies into a common class, unless we are guaranteed the mapping represents true equivalence. This is illustrated in figure 1, which shows mappings between classes in three vocabulary sources.

We have devised and implemented an algorithm called kBOOM that assists a biocurator in merging ontologies. The kBOOM algorithm takes as input a *probabilistic* ontology which consists of a combination of logical axioms and hypothetical axioms, and returns the most likely ground ontology. Hypothetical axioms can be generated using a variety of methods, including using predefined curated mappings. The resulting ontology will include only precise logical axioms connecting classes, such as SubClass (⊑) and Equivalence (≡) axioms. Any classes connected via equivalence axioms can be safely merged (i.e. merging produces a structure with the same logical entailments)

**Fig. 1.**
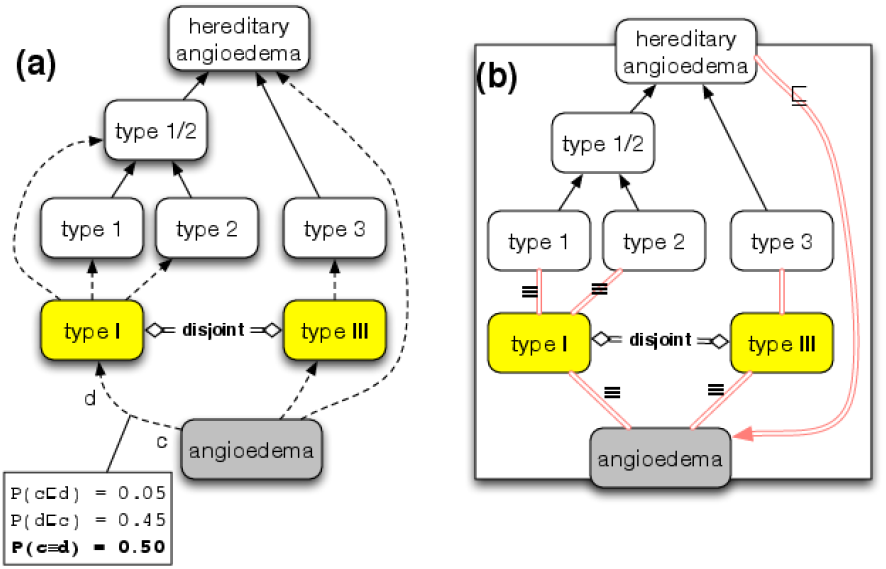
(a) Mappings between 3 vocabularies (shown in different colors). Logical relationships (axioms) within an ontology (subsumption) shown in solid lines, Arrows denote SubClassOf (e.g. *type-1* ⊑ *type-1/2*). In the yellow ontology, types 1 and 3 are declared *Disjoint* (i.e. no shared subclasses). Dashed lines represent loose mappings (b) one possible configuration with hypothetical axioms derived from mappings shown as double red lines. In this example, all mappings interpreted as equivalence (≡) except *hereditary angioedema* ⊑ *angioedema* (white to grey). Note that due to the logical properties of ⊑ and ≡ all nodes in the enclosing box are inferred to be in a mutually equivalent clique, yielding incoherency (via disjointness axiom).

We are applying kBOOM to merge a mixture of ontologies, databases and vocabularies which describe diseases or disorders, into the Monarch Disease Ontology (MonDO), because no one single source provides a comprehensive picture of the disease space. For example, the Disease Ontology (DO)[3] provides a broad classification of disease types, but does not enumerate all the diseases in Online Mendelian Inheritance in Man (OMIM)[1].

Conversely, OMIM provides largely a flat list without a hierarchy, and typically only covers disorders with genetic etiology. ORDO[8] is derived from Orphanet and has an emphasis on rare disease, and employs different classification criteria from DO. When previous attempts have been made to unify these, the results are frequently incomplete. For example, MedGen[2] includes OMIM and Orphanet, but many of the entries, such as that for *Hyperekplexia hereditary* lack a classification. Here we explore an ontology merging strategy as one possible solution to this problem.

## 2 METHODS

Our approach takes a collection of ontologies *O*_1_,…,*O_υ_*, and produces a coherent well-connected merged ontology *O^M^*, in which the classes are connected using OWL logical axioms. We include vocabularies and databases in our definition of ontology. The input may also include a set of previously curated mappings *M* ∈ *C* × *C*, where *C* is the total set of classes in all sources. We assume that a mapping means that the two classes have some substantial degree of similarity, but do not assume any logical semantics.

Our approach consists of a pipeline with two steps:

1. *Generation of a probabilistic ontology with prior probabilities*. Prior probabilities can be generated using a variety of methods, such as lexical approaches, or by using existing curated mappings.
2. *Estimation ofthe most likely ontology*. We attempt to maximize the posterior probability for different combinations of axioms, utilizing the complete OWL semantics^1^. Once this has been determined, additional post-processing can be performed, such as merging equivalent classes.

We focus on this second step as it is a generalizable method, and the primary contribution of this paper. We first describe the *general* approach which is applicable to OWL ontologies of any expressivity, and we then describe a *specific* approach which can be applied to simpler ontologies in which the probabilistic aspect utilizes only concept inclusion and equivalence axioms.

To illustrate the first step with a concrete example, we give an example of a procedure for estimating prior weights for disease ontology mappings, and outline how we are using the combined procedure to build the Monarch merged disease ontology (MonDO).

### 2.1 Ontology structure inference: general approach

#### 2.1.1 Probabilistic Ontologies

The input to the structure inference procedure is a *probabilistic ontology*:

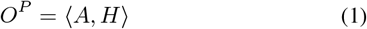

where *A* consists of *logical axioms A*_1_,…,*A_ι_*, and *H* consists of *hypothetical axioms H*_1_,…, *H_n_*.

Both of these axiom sets involve any combination of classes from the union of sources, which we denote *C*. However, a common use case is that *A* consists purely of within-source axioms (e.g. two independent disease ontologies) and *H* consists of hypothetical axioms derived from cross-ontology mappings.

The joint probability distribution (JPD) is:

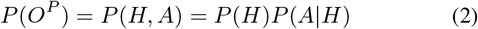

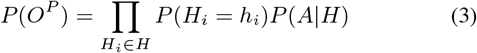

where *h_i_* is a boolean representing the truth value of hypothetical axiom *H_i_*

For *P*(*A|H*) we assume a uniform probability, except in the case where the ontology *O^P^* is *incoherent*, i.e. there exists a class *c* ∈ *C* such that c is *unsatisfiable*, where *unsat*(*c*) = *O^P^* ⊨ ≡⊥. This situation can arise whenever the ontology violates any logical constraints, encoded in OWL. This state can be checked using a standard deductive OWL reasoner.

We attempt to find the most likely ontology, i.e. set of values *h*_1_,…, *h_n_* that maximizes the posterior probability.

As there are 2^|*H*|^ combinations, we use a number of techniques to reduce the search space. For the first of these, we *factorize* the calculation by splitting the ontology into sub-modules.

#### 2.1.2 Factorization via module extraction

A *modularization* procedure takes an ontology *O* and produces *N* ontology modules. We can apply the analogous procedure to probabilistic ontologies and produce *k* probabilistic modules. A number of different modularization strategies can be applied. For some problems it may be possible to produce *k* modules that are independent, in which case the probability can be factorized as:

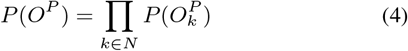

In other cases, modules may have dependencies; we do not cover this here, approaches may involve Bayes networks or factor graphs.

#### 2.1.3 Greedy selection search procedure

Even after factorization, the number of possibilities may still be prohibitive (a module size of 100 nodes is not atypical, which has 4^1^00 configurations). In order to further reduce the search space, we apply a greedy approach. Here, when searching for solutions for any module *O_k_* we iteratively take the axiom for *H_k_* with the highest probability and assume it to be true. We repeat this until the number of hypothetical axioms is reduced to a user-specified threshold *T*.

We immediately reject any axioms that lead to *O^P^* becoming incoherent. We attempt to re-modularize at each stem: the selection procedure may lead to the current module being splittable into sub-modules.

### 2.2 Ontology structure inference: Approach for *AL*^−^-ontologies

We have implemented a procedure for inferring ontology structures where the hypothetical axioms consist only of the ⊑ (SubClass) or ≡ (Equivalence) constructs, applied between two named classes *C_i_, C_j_* ∈ *C*. Formally:

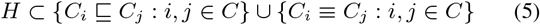

This is a subset of the Description Language AL (Attributive Language), which we denote *AL*^−^. Note we do not restrict the expressivity of the non-hypothetical logical axioms *A*.

Limiting the expressivity of the probabilistic subset reduces the search space, as there are fewer possibilities. For any pair of classes *c,d* in a mapping, there are only four possibilities: *c* ⊏ *d, d* ⊏ *c, c* ≡ *d, c* ⋢ *d* ∧ *d* ⋢ *c*. This limitation also allows for the extraction of smaller modules.

For the ontology mapping use case, we also assume an additional logical constraint in the form of *AllDistinguishableFrom* axioms. Here we assign each class in the ontology to a *source*, and we constrain such that there are no equivalent pairs of classes within a source.

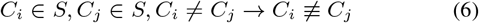

#### 2.2.1 Extracting Locally Independent Modules for AL^−^

By limiting the probabilistic subset to only SubClass and Equivalence axioms, we can exhaustively partition the ontology into independent submodules, based on whether the probability of any two members of the partition being equivalent is non-zero.

Here we split the ontology into *k* modules such that each module is independent. We take all hypothetical equivalence axioms in *H* and assume *P*(*H_i_* = *true*) = 1, thus creating a *maximally equivalent* ontology *O’* (see figure 1b for example). We then obtain the set of all entailed equivalence cliques *N* from *O*’ (using any reasoner that is complete for the profile of *A*). For this step, we ignore logical axioms such as disjointness axioms that can lead to incoherency.

We then iterate through all cliques *N*_1_,…, *N_n_* ∈ *N* and extract a probabilistic module *O^N_i_^*, where the module contains all classes in *N_i_*, plus all logical axioms in *A* that reference any class in *N_i_*, plus all hypothetical axioms in *H* that connect a class in *N_i_*.

This corresponds to the factorization in Equation 4.

#### 2.2.2 Estimating prior edge probabilities for disease mappings

kBOOM takes as inputs hypothetical axioms with assigned prior probabilities. For this, we make use of existing mappings provided by source databases and ontologies. These are manually curated and are generally of high quality, in that a mapping typically corresponds to a biological or clinical connection. However, they are often incomplete, and do not have a strict meaning, ranging from equivalence, subclass, superclass, without any guarantees on the interpretation.

We turn each mapping into a hypothetical axiom and use a variety of heuristics and rules to generate weights for different axiom types. Given a pair of classes *c* and *d*, we have rules such as:

1. *c* ≡ *d* +++ if *label contains ‘type’ or ‘complementation’*
2. *c* ≡ *d* ++ if *StrictLexicalMatch(c,d)*
3. *c* ≡ *d* + if *NonStrictLexicalMatch(c,d)*
4. *c* ⊑ *d* ++ if *c* ∈ `OMIM`, *d* ∈ `OrdoGroupOfDiseases`
5. *c* ⊑ *d* ++ if *c* ∈ `OMIM` and *d has a subclass*
6. *c* ⊑ *d* + if *c has a NarrowSynonym = label(d)*
7. *d* ⊑ *c* + if *c has a BroadSynonym = label(d)*
8. *c* ⊑ *d* + if *label(c) SubStringOf label(d)*

The number of +s denotes the weighting (we omit the actual calculations and numbers here for brevity, code on GitHub for details). For example, if the labels or exact synonyms match exactly, and the string is of a form like *Foo type 2*, then our confidence in an equivalence match is higher. In addition to lexical criteria, we use ontology structure criteria. For example, Orphanet classifies diseases into meta-classes such as GroupOfDisorders and EtiologicalSubtype, with the latter always being a superclass of the former - these can be used to weight our confidence of equivalence vs subclass matches.

We can also look at the distribution of mappings between two sources. In general, if a single class in *S*_1_ is mapped to multiple classes in *S*_2_, then the relationship between the two classes is more likely to be a superclass relationship, because in general classes have more children than parents. We make an exception for Orphanet, which frequently groups disorders under multiple classes on a per-phenotype basis.

The rules above do not exhaustively cover all cases, and a number of additional lexical normalization steps are applied. These are not described here for brevity.

## 3 RESULTS AND DISCUSSION

Our Java implementation of kBOOM is in GitHub^3^, and currently only implements the special case for *AL*^−^.

We created a GitHub repository for the disease use case^4^ containing both the rules to generate hypothetical axioms from pre-defined mappings and the generated ontology. Our inputs are DO, OMIM labels, Orphanet, MeSH, MEDIC, OMIA, GARD and DECIPHER (with some only contributing labels). The largest cluster we have observed has 184 classes, constructed from a highly connected set of mappings for *retinitis pigmentosa, code-rod dystrophy* and *Leber Congenital Amaurosis* subtypes. Over 65k hypothetical axioms were generated from mappings, of which 11,800 were interpreted as equivalence axioms, allowing the safe merging of multiple duplicate classes across ontologies.

Our next steps are to perform a full evaluation of these results. Figure 2 shows an example of how a module has been resolved.

**Fig. 2.**
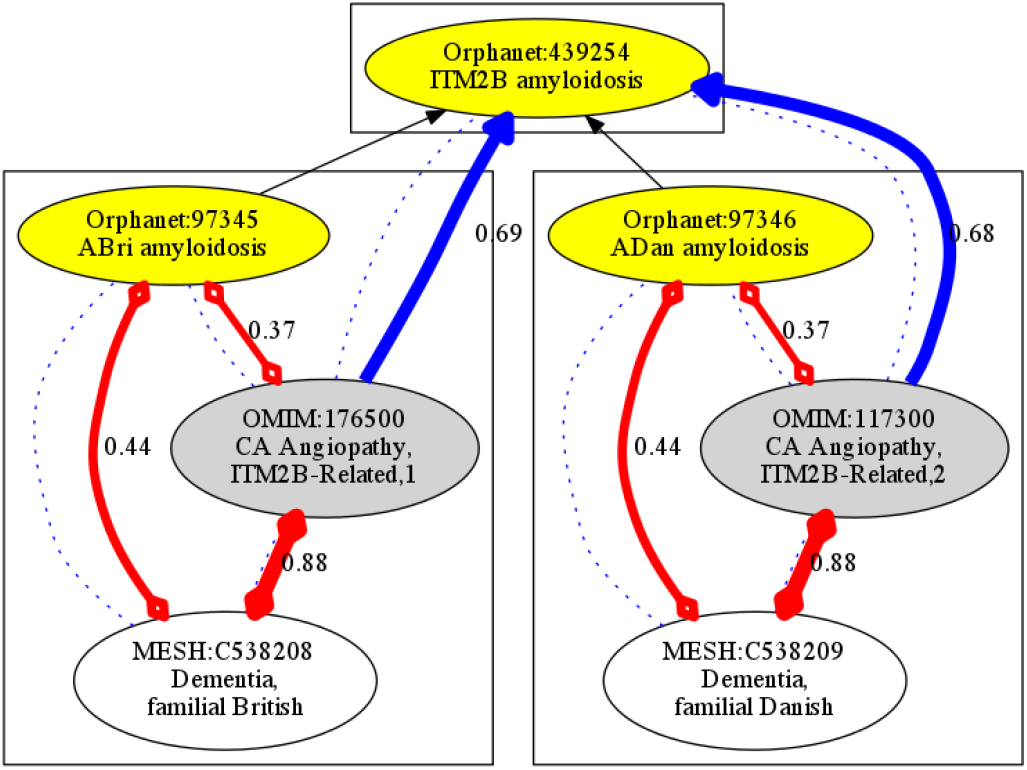
Module resolution graph exported by kBOOM; Initial input is nodes plus solid arrows (SubClassOf axioms in ORDO). Dotted lines are supplied mappings (no logical interpretation). Figure shows inferred most likely configuration. equivalence=red, subclass=blue, with prior probabilities written as edge labels (thick lines more probable). Enclosing boxes denote equivalence cliques, which can be merged to a single class, yielding a grouping class with two children.

### 3.1 Tolerance and detection of mapping errors

When we assign prior probabilities, we assume a low error rate in source-provided mappings, and thus for larger modules these are rarely rejected, even if it leads to an overall more probable structure (due to the greedy selection procedure). As a pre-processing step we take large modules and iteratively test removal of axioms to determine if the module can be broken into semantically distinct submodules, which can sometimes detect incorrect mappings (e.g. ^5^). We plan to fold this into the probabilistic approach.

### 3.2 Integration with curation

One advantage of our approach is that it fits well with both manual curation and knowledge-driven approaches. The more knowledge provided by a curator, the better the results; at the same time, the approach is robust to incomplete information, attempting to estimate the most likely structure, and always producing a coherent result, if it can be found.

A curator can provide different kinds of information, and this can be done iteratively. The most basic information to be provided is a manually vetted curation. Additional information can be in the form of prior probabilities for different axiom types; for example, if a curator strongly believes a mapping denotes equivalence, this can be provided (and if the curator is highly confident, the axiom can be removed from the probabilistic set and added directly into the ontology).

The search procedure uses an OWL reasoner, so other axiom types can be freely used; e.g the curator can provide defining equivalence axioms to class expressions. The curator can also provide constraints on possible models using constructs such as disjointness axioms, or taxon constraints or some of the other logical constraints developed in GO[6] and Uberon.

### 3.3 Comparison with other approaches

There are a number of probabilistic approaches to ontology mapping, but most are aimed at generating rather than interpreting mapping. One exception is the OMEN algorithm[5] which is similar to our approach in that it generates probability distributions for axiom types based on mappings, and factorizes the joint probability distribution into a bayesian network. However, the approach has certain assumptions that are problematic for the disease mapping use cases, such as the combined ontology containing no cycles. We aim to do further investigation into a generalized approach that unifies OMEN and kBOOM.

In order to fully evaluate our method, we plan to compare the disease merging results with other combined disease resources such as MedGen and EFO[4]. Additionally, we apply on other domains such as anatomy and compare to gold standards such as Uberon.

### 3.4 Future Plans

#### 3.4.1 Tree Search Algorithm (POETS)

Here we propose a tree search approach, as an alternative way to search the space of 2^|^*H*| possible solutions. This can either be used on *O^P^* directly, or in combination with a modularization approach. We call this POETS (Probabilistic Ontology Entailment Tree Search).

Each combination of hypothetical axioms constitutes a ‘possible world’. We represent each possible world as a node in a tree. The root node of the tree is the empty set, i.e. no hypothetical axioms are assumed either true or false. We greedily explore outwards from each node, assuming a new hypothetical axiom each time. At each node we will compute the entailed logical axioms, and use them to calculate posterior probability.

Search is terminated after some amount of time, and the most likely leaf node is selected (if there are no leaf nodes with *P* > 0 then there is the option of selecting further or simply returning incomplete results).

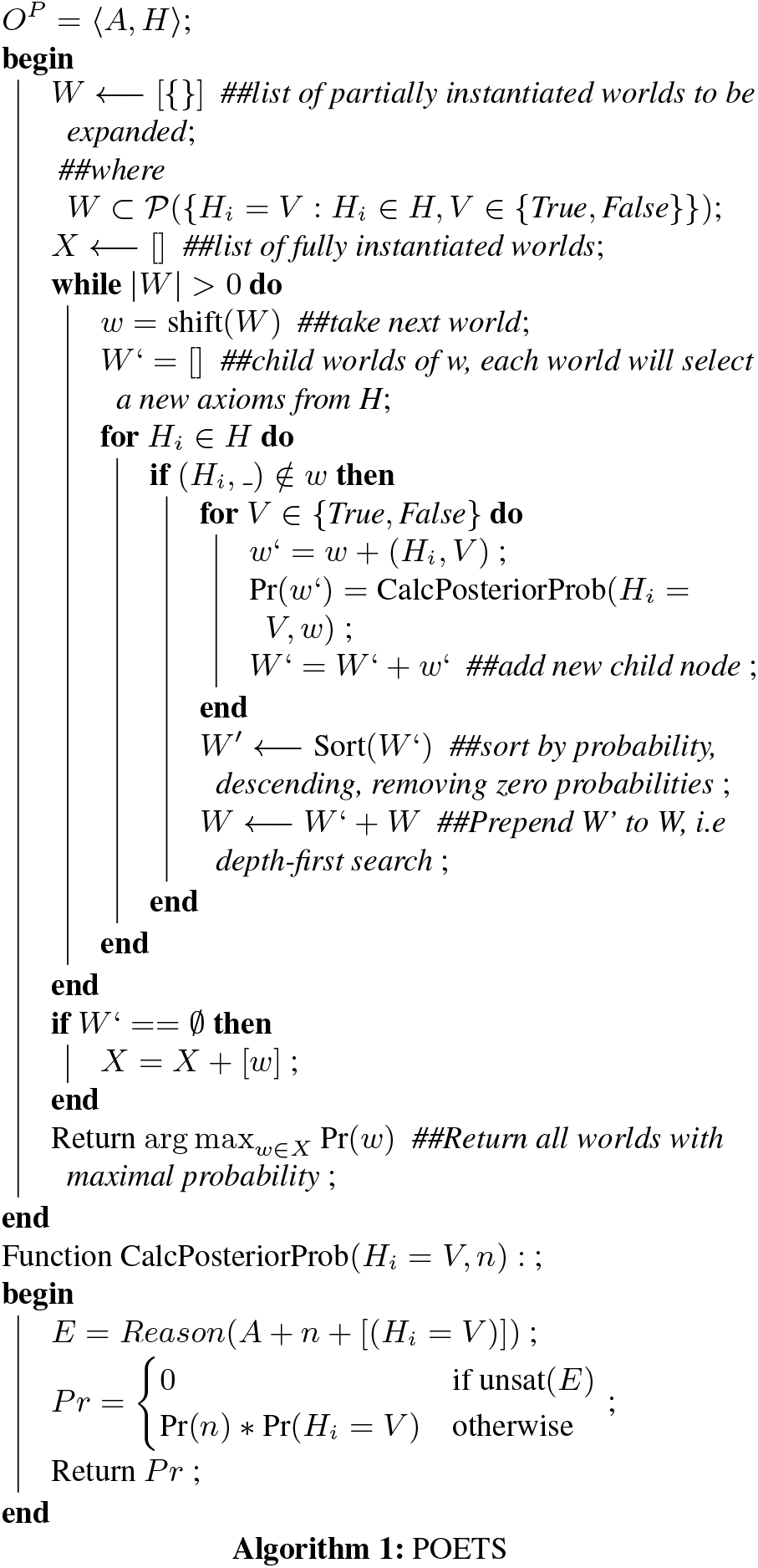

The base tree-based search algorithm is defined in algorithm 1. The intent is to implement this using a reasoner that utilizes *immutable* in-memory models of ontologies and inference, since each child node can incrementally reasoner over the parent world.

## Footnotes

1 https://www.w3.org/TR/owl2-direct-semantics/

3 https://github.com/monarch-initiative/bayes-owl-ontology-merging

4 https://github.com/monarch-initiative/monarch-disease-ontology/

5 https://github.com/DiseaseOntology/HumanDiseaseOntology/issues/134

